# GWA-X: An Extensible GPU Accelerated Package for Permutation Testing in Genome-Wide Association Studies

**DOI:** 10.1101/2024.09.15.613119

**Authors:** Mulya Agung, Albert Tenesa

**Affiliations:** University of Edinburgh

## Abstract

Genome-wide association studies (GWAS) aim to identify genetic variants that are associated with a trait or disease. The scale of genomic datasets has increased to millions of genetic variants and hundreds of thousands of individuals, opening the possibilities for GWAS discoveries. However, large-scale GWAS analyses are prone to high false positive rates because of the multiple testing problem. Permutation testing is the gold standard for maintaining false positive rates, yet it is impractical for large-scale GWAS because it demands vast computational resources. This paper presents GWA-X, a software package that can exploit GPU potential to accelerate permutation testing in GWAS. Unlike previous methods, GWA-X employs a novel whole-genome regression method and a permutation testing strategy to batch the computations of many genetic markers. It achieved a two-order magnitude speed-up compared with the existing CPU-based and GPU-based permutation methods. In addition, it provides an extensible package for GWAS permutation testing on GPUs.

## 1 Introduction

Genome-wide association studies (GWAS) scan a genome-wide set of genetic variants or markers in a group of individuals to identify variants associated with a trait or disease. The growth of GWAS over the past decade has identified numerous associations for complex human traits and diseases. In recent years, the scale of genomic datasets has increased to millions of genetic variants and hundreds of thousands of individuals, opening the possibility for discoveries from GWAS and omics studies [1, 2].

Despite the availability of large genomic data, large-scale GWAS demands vast computational resources due to the many variants that need to be tested [3, 4]. In addition, conducting GWAS with millions of genetic variants has a severe problem of the potentially high number of false positive results due to many variants tested [5, 6] and non-normal phenotypic distributions [7, 8].

Permutation tests are the gold standard for multiple testing correction in GWAS because they can reduce false positives while maintaining statistical power [9–11]. However, permutation tests are computationally demanding because they require many statistical tests to obtain a statistical distribution. The growth of scale in genomic datasets further increases the computational burden of permutation tests. The scale is expected to grow to billions of variants and millions of people in the future [1, 12]. Consequently, permutation testing is hardly practical in large-scale GWAS.

The end of Dennard scaling and the imminent end of Moore’s law have led to the necessity of using specialized computing processors such as graphics processing units (GPUs) to improve computational efficiency [13–15]. Because GPUs can perform numerical calculations faster than central processing units (CPUs) at comparable energy costs, they could significantly reduce GWAS’s computational time and energy consumption. Although GPUs have been widely used in AI research [16], the use of GPUs in GWAS is still limited because existing GWAS tools that work well for CPUs cannot take advantage of GPUs due to hardware GPU constraints that are not present in CPUs. In addition, not all computations in GWAS permutation testing are suitable for GPUs. A collaboration between CPU and GPU is necessary to achieve high computational efficiency [17].

Several linear algebra libraries have been developed for GPUs [18, 19]. However, these libraries cannot efficiently parallelize GWAS’ regressions because they aim to compute a regression of a large matrix problem in parallel to use all the GPU cores [20]. The regression in GWAS is computed for each variant or a group of variants, resulting in many regression computations of small matrices instead of one large matrix. Thus, these libraries cannot efficiently utilize GPUs and cannot achieve significant performance improvements compared to CPUs.

Besides the computational demand, diverse permutation testing approaches have been developed for different applications in GWAS [10, 21, 22] simultaneously with novel regression methods to account for the population structure in the emerging genomic data [3, 23]. Meanwhile, the growth of genomic data size has imposed new computing system requirements, such as biobank platforms [2, 24]. GWAS permutation testing could benefit from the latest methods and computing systems. However, existing GWAS and permutation testing tools cannot be easily extended due to their architecture limitations. Therefore, there is a demand for an extensible package for GWAS permutation testing that can adapt to new developments.

This paper presents GWA-X, a software package that leverages GPUs to accelerate GWAS permutation testing. In contrast to previous methods, GWA-X batches regression computations of many genetic variants to achieve substantial acceleration on GPUs. The **contributions** of this paper are as follows.

- A whole-genome regression method for GPUs while accounting for population structure, including a genetic marker-based batch computation algorithm and a chunked data processing technique to handle large genotype data and use multiple GPUs.
- A permutation testing strategy for GPUs, including a memoization technique and a CPU-GPU collaboration strategy to accelerate permutation tests.
- An extensible and scalable software package that allows the addition of new regression models, permutation testing approaches, and computing system requirements.

## 2 Related Work

Several libraries have been developed to accelerate batch matrix computations on GPUs [19, 20, 25], which mostly focus on square matrix problems or independent tall-skinny matrices. However, a GWAS analysis requires regressions of many matrices of genetic markers and the same matrices of the trait and covariates. These libraries cannot parallelize many regressions enough to maximize GPU occupancy. Hence, the computational performance of these libraries will not scale with many genetic markers in GWAS.

Most of the existing GWAS permutation testing methods focused on CPUs and cannot use GPUs [9, 26–28]. More recent methods use GPUs to accelerate the permutation tests in GWAS [10, 29]. However, these methods rely on existing GPU libraries that cannot fully utilize GPU’s potential performance. In addition, the previous GPU-based methods cannot handle datasets larger than GPU memory and are not extensible to different permutation testing approaches and computing system requirements.

In contrast with the GPU libraries and the previous methods, GWA-X batches regression computations of many genetic markers and uses a chunked data processing technique to handle large datasets. The evaluation of this paper demonstrates the superiority of GWA-X over the previous CPU-based and GPU-based methods and an implementation using the current state-of-the-art GPU library.

## 3 Method

Typical GWAS approaches perform generalized linear models (GLMs) to test associations of genetic variants or markers and traits depending on whether a continuous or binary trait is analyzed [5]. For each genetic marker, a GLM regression is given by the following equation:

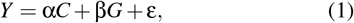

where *Y* is a trait or disease of interest, *G* is a genetic marker, *C* represents a matrix of covariates, and ε is the residual.

To account for sample relatedness or population structure, an additional step is required to reduce the bias caused by genetic relatedness among individuals, such as in linear mixed models (LMMs) [4, 11]. However, LMM requires the computation of the genetic relationship matrix (GRM), which is computationally expensive with large samples [30]. Alternatively, ridge regression is a machine learning approach that can estimate the sample relatedness and is more computationally efficient [3, 31]. In addition, it allows population structure correction to be decoupled from association testing and permutation tests.

GWA-X adopts the previous method [3] to compute the genetic predictors correlated to the trait,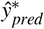, that estimates the sample relatedness. The predictors are obtained by fitting the model in Equation (2) using ridge regression across a range of shrinkage parameters, where *W* is a matrix derived from *G* with substantially fewer columns, and η is estimated as 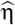 by adding an *L*2 penalty to the log-likelihood function of the model in Equation (2). After 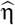 has been estimated, the predictors can be obtained as in Equation (3), using the leave-one-chromosome-out (LOCO) scheme by ignoring columns of the chromosome containing the genetic marker being tested when calculating the predictors. Then, instead of using the original trait values (*Y*) for Equation (1), GWA-X uses the corrected trait values (*Ŷ*) obtained from removing the predictors from *Y* as in Equation (4). More details on the ridge regression method can be found in the original paper [3].

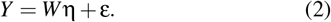

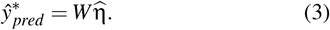

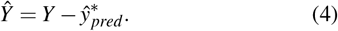

In GWAS permutation testing, the trait values are permuted *Q* times while keeping the variants intact to account for linkage disequilibrium (LD), and the statistical tests are repeated for each permuted dataset [9, 22, 27, 29]. The test results are then used to generate the distribution under the null hypothesis of no association. Statistical significance (*p*-value) of the permutation testing (*P*_*perm*_) can be calculated by Equation (5), which is the fraction of permuted datasets for which the value of the test statistic (*T*_*q*_) is equal or greater to the value observed on the original data (*T*_*obs*_).

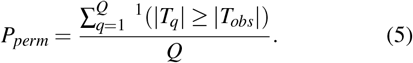

Accordingly, the number of regression computations will increase linearly with the number of genetic markers and permutations. Regressions with many variants and large sample sizes are computationally demanding even without permutation [3, 4]. Therefore, accelerating the regressions is key to achieving a high computational efficiency in permutation tests.

GWA-X accelerates the computational performance of GWAS permutation testing in two steps: it increases the degree of parallelization for regression computations on GPUs. Then, it employs a CPU-GPU collaboration strategy to accelerate permutation tests. The following subsections detail these steps.

### 3.1 Massively Parallel Regression on GPUs

GPUs have two memory types: on-chip and global memories. On-chip memory is faster but much smaller than global memory. In GPU programming, computation tasks are executed as units, called threads, organized into a grid of thread blocks. Each thread block can share data via an on-chip memory space known as shared memory, which is typically small and can only be accessed by threads within the same block. Before the computation, threads load the data from the global memory to the on-chip memory. Unlike CPUs, adjacent threads must access adjacent data elements in global memory to achieve high speeds on GPUs. That is why methods designed for CPUs often perform poorly on GPUs [32].

GPUs have thousands of processing units that can carry out many calculations in parallel, which is vital to accelerating regression computations in GWAS. A regression method must maximize the number of processing units used in parallel to achieve high computational efficiency, whereas memory access is slower than computation. GWA-X increases the memory access throughput from/to GPU’s global memory by coalesced access to maximize the use of GPU’s memory bandwidth [20, 32].

Another key aspect is the organization of threads because it determines how many computation tasks can run in parallel and share data through shared memory. GWA-X uses the grid shown in Figure 1, which allows different markers and samples to be computed in parallel. Simultaneously, to use the fast shared memory, the threads of the same block compute the regression for the same marker. For example, regression stores temporary data on shared memory for matrix multiplication to reduce the number of accesses to global memory.

**Figure 1:**
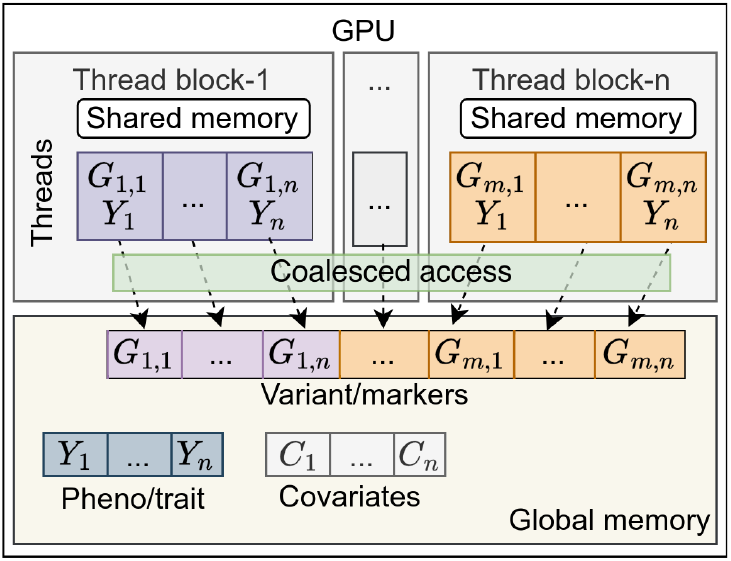
GWA-X’s thread organization on GPU

In GLM, the genetic and fixed effects (β, α) can be calculated using Equation (6), where *X* is a concatenated matrix consisting of a genotype vector with a covariate matrix (*G*| *C*) and *Y* or *Ŷ* as the vector of trait values. Thus, the regression requires an matrix inverse operation, which is computationally intensive. GWA-X accelerates regressions by computing matrix inverse of many markers in parallel. It uses an algorithm extended from the previous parallel Gauss-Jordan algorithm [33]. The extended algorithm computes the matrix inverse of different genetic markers on different thread blocks, and the matrix inverse of a genetic marker is computed by all the threads of the same thread block to use shared memory.

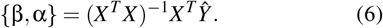

GPUs have limited memory capacity, and the data required might not fit entirely into the GPU memory, such as in wholegenome regression. GWA-X addresses this problem by chunking genotype data for data transfers and processing, including decompressing compressed genotypes on GPUs to increase GPU occupancy. The chunk or block size can be specified by users according to the GPU memory size. As sequential data transfers to process all the chunks can delay GPU computations, GWA-X reduces the delay by overlapping the data transfers with GPU computations (Figure 2). In a system with multiple GPUs, the genotype chunks are distributed to the GPUs, and regressions are computed in parallel on multiple GPUs (Figure 3).

**Figure 2:**
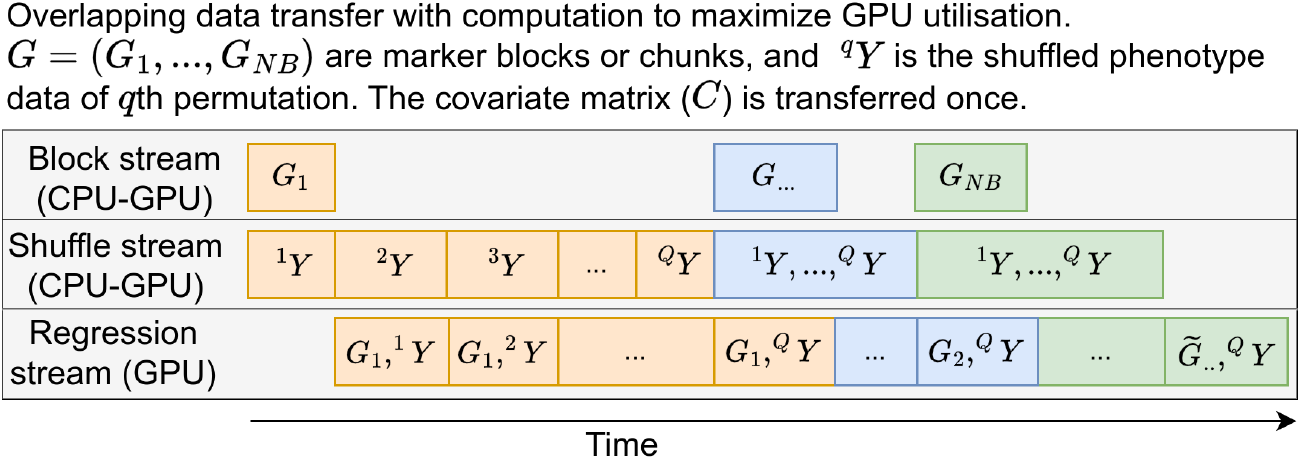
Overlapped computations with data transfers

**Figure 3:**
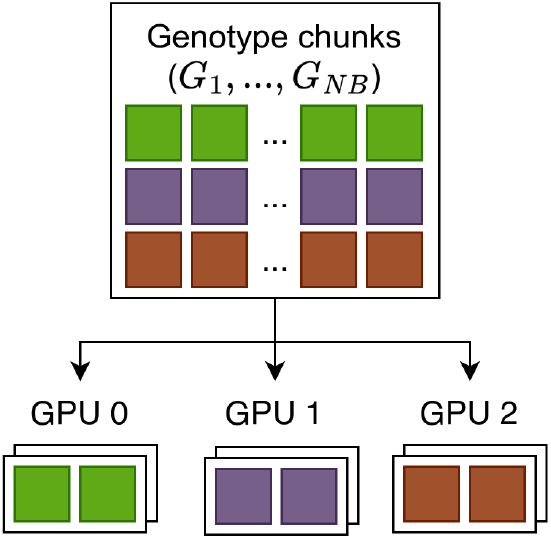
Genotype chunk distribution on multiple GPUs

### 3.2 Permutation Testing Strategy

CPUs and GPUs have unique features and strengths. Hence, dividing computational workloads among them can significantly improve computational efficiency [17]. For instance, data shuffling involves much more memory operations than numerical calculations. Hence, permuting trait values of large samples can be slower on GPUs than on CPUs due to the GPU’s memory access latencies. In addition, transferring the permuted data from/to CPUs and GPUs sequentially can delay GPU computations. GWA-X shuffles the trait data on CPUs and transfers them with chunked genotype data to GPUs overlapped with regression computations, as shown in Figure 2.

In permutation tests, trait values (*Y* or *Ŷ*) are shuffled for each permutation *q*, denoted as (^*q*^*Ŷ*). Then, the β and α coefficients are calculated for each permutation (β_*q*_, α_*q*_) using Equation (7). The permutation test statistic (*T*_*q*_) for a genetic marker is calculated using Equation (8), where β_*q*_ and *SE*_*q*_ are the coefficient and standard error of the permutation *q*, respectively. Therefore, the genotype covariance matrix, (*X*^*T*^ *X*)^*−*1^, needs to be computed only once and can be reused for the whole permutation tests, including for the *SE* calculation.

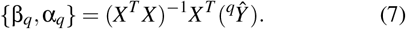

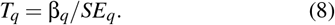

Instead of computing standard errors (*SE*s) sequentially for each marker, GWA-X calculates them for a batch of markers at once on GPUs. For each marker, *SE*s are calculated using the following matrix algebra:

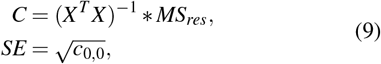

where *C* is the parameter covariance matrix, *c*_0,0_ is the value from *C*’s diagonal vector that corresponds to β, and *MS*_*res*_ is the mean square of residuals for each permuted dataset. GWA-X applies a memoization technique that stores and reuses the covariance matrices of genetic markers to reduce computation time.

## 4 GWA-X Architecture

GWA-X’s architecture, shown in Figure 4, comprises (1) a system layer containing computing system backends, (2) a library layer implementing regression models and permutation testing approaches, (3) and an application layer to integrate the library and system layers.

**Figure 4:**
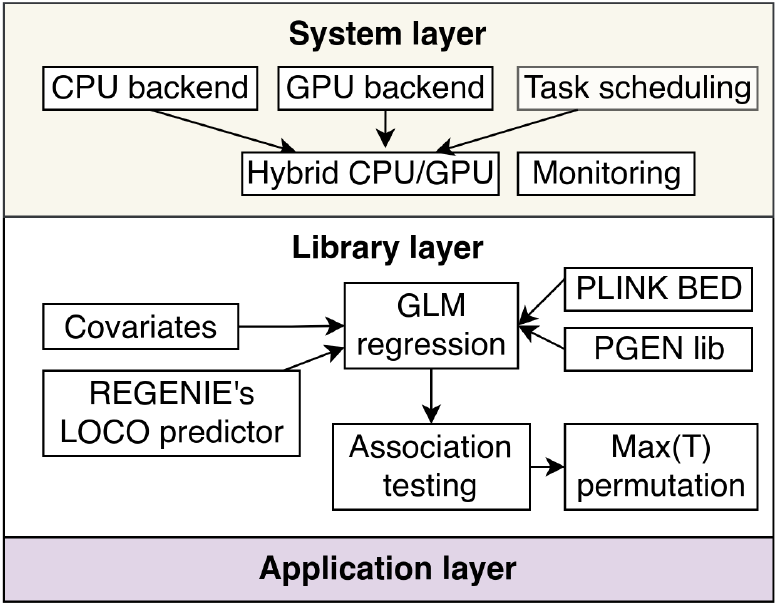
The software architecture of GWA-X

As the size of genomic data will increase in the future [2], the use of distributed computing systems is necessary. Accordingly, the permutation methods must be scalable to distributed CPU-GPU systems, imposing new system requirements. The system layer allows additional backends to use different systems, such as multiple GPUs and distributed systems, without changing the library layer. This flexibility also broadens GWA-X applicability to various computing systems, such as biobank platforms [2, 24]. Besides system backends, the system layer provides a task scheduler to optimize parallel executions on CPUs and GPUs and a monitoring subsystem to track computational performance and energy usage.

The library layer contains implementations of regression algorithms and permutation testing approaches with a programming interface to reuse them. It presents the advantage that the regression and permutation testing routines can also be used for different analyses to benefit from GPUs, such as multi-omics [34] and rare variant analysis [22]. The application layer integrates the library and system components and provides the user interface for conducting permutation tests.

### 4.1 Implementation

GWA-X is an active open-source project^†^ developed at the University of Edinburgh, UK. It fully integrates with the Python environment and is easy to install using the Pip package manager. The regression method is built from scratch and compiled into machine code using the Numba compiler [35] to achieve comparable performance to a compiled programming language.

At the time of writing, a whole-genome regression method for quantitative traits and a standard permutation testing approach have been implemented. For accounting for population structure, GWA-X can use the LOCO prediction file generated from Step 1 of the REGENIE tool [3] and a matrix of covariates. It supports the common PLINK BED and PGEN genotype formats [28] and CSV formatted data of phenotype or trait.

As a future roadmap, GWA-X will support logistic regression for binary traits, adaptive permutation [21], and system backends to use distributed CPU-GPU systems. GWA-X currently requires REGENIE for computing ridge regression to generate the genetic predictors. Thus, another future work is extending GWA-X to support ridge regression on GPUs. The extensible architecture will allow future development to be applied modularly, requiring no or minimal modification to the implemented parts and different layers.

## 5 Data and Simulation

The statistical and computational performances of GWA-X are evaluated on simulated genotype-trait datasets. To mimic the linkage disequilibrium structure of real datasets, the genotype data were simulated from the HapMap3 (release 2 build 36) haplotype data using HAPGEN2 [36], resulting in up to 430,000 samples and 1,002,365 array SNPs with minor allele count higher than 5.

Artificial traits were simulated for 5,000 and 10,000 causal SNPs as causal markers from the genotype simulated from HapMap3. The effect sizes for the causal SNPs (β) were sampled from a normal distribution with a mean of zero, where the variance was determined based on the desired proportion of trait variance explained by the causal SNPs, *h*^2^. The effect from population structure (α) was set so that the proportion of the trait variance explained by the top principal component was 10%. Random noise (ε) was generated from a normal, Student’s t, or gamma distribution. The genetic predictors for the simulated traits were generated using REGENIE.

## 6 Evaluation

The evaluation comprises computational efficiency and statistical performance comparisons on the simulated datasets.

### 6.1 Computational Efficiency

A relatively recent method, permGWAS [10], has been proposed for accelerating permutation testing on GPUs. In contrast to GWA-X, permGWAS uses the PyTorch library [19] to perform batch-wise matrix computations on GPUs. Figure 5 shows the computation times of both methods on an NVIDIA A5000 GPU card with a simulated dataset of 4,000 samples and one million genetic markers, a quantitative trait, and a matrix of covariates. GWA-X outperforms permGWAS and achieved more than two orders of magnitude speed-up in permutation tests (129.9x on average), reducing the computational time from 3.4 hours to 74 seconds. The evaluation could not run permGWAS on larger samples because it requires a large memory space to load whole genotype data into memory. This fact shows the necessity of the data chunking technique in GWA-X to allow permutation tests with data sizes larger than CPU and GPU memories.

**Figure 5:**
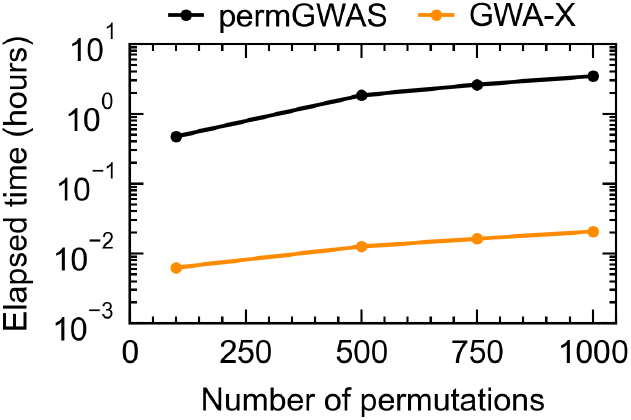
The computational performance of GWA-X and permGWAS with 4,000 samples and one million markers

To investigate computational efficiency on larger samples, the evaluation compares the computational performance and energy consumption of GWA-X to a CPU-optimized software, PLINK [28], and an implementation using the GPU-accelerated PyTorch [19] library (PyTorch-impl), which uses the torch.linalg module to perform batch matrix computation on GPUs. In contrast with permGWAS, PyTorch-impl uses genotype data chunking to handle large samples. PyTorch-impl is available in the GWA-X project repository. The evaluation uses the simulated dataset of a quantitative trait, a covariate matrix, and a genotypes with 430,000 samples and up to 500,000 genetic markers.

GWA-X and PyTorch-impl were run on an NVIDIA A5000 GPU card, while PLINK was run with the -linear option on a 16-core Intel Xeon 5128 CPU @2.3 GHz. For permutation tests, GWA-X performance is compared with PLINK (-linear mperm) on a single node of the ARCHER2 super-computer [37], equipped with 128-core AMD EPYC 7742 processors.

Figure 6 shows the computational performance of regression. GWA-X achieved a faster regression by a hundredfold compared with PLINK (95.5x on average) using only one GPU card, more than tenfold faster than the PyTorch-based implementation (12.5x on average). Figure 7 shows the energy consumptions and carbon emissions of GWA-X and PyTorch-impl, measured using CodeCarbon [38] that is implemented in GWA-X’s system layer. GWA-X reduced energy and CO2 emissions by 88.4% from faster regressions.

**Figure 6:**
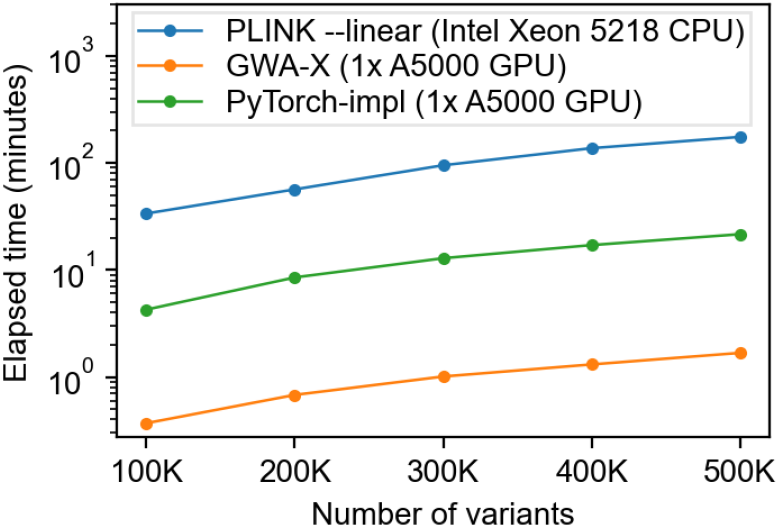
The computational performance of regression for quantitative trait analysis

**Figure 7:**
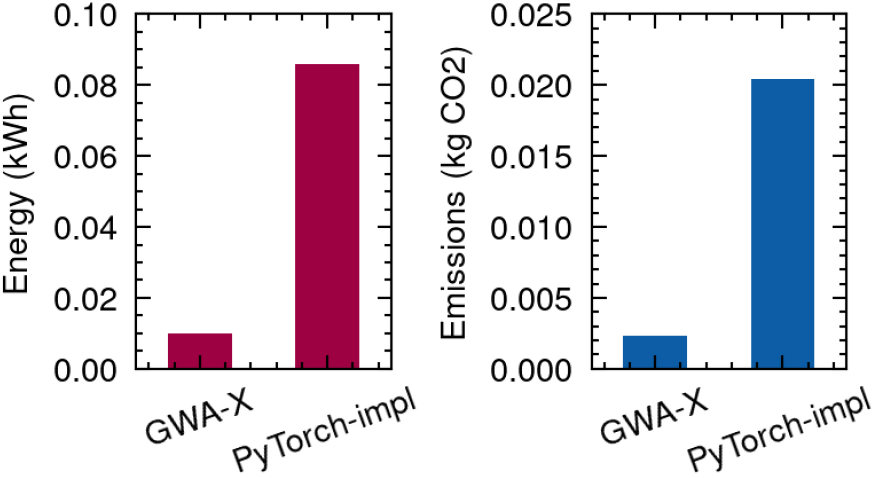
The estimated energy usage and carbon emission of GWAS regression

For permutation tests, GWA-X achieved a speed-up ratio of more than two orders of magnitude (99.5x on average) compared with PLINK on ARCHER2, as shown in Figure The results suggest that GWA-X can save substantial cost and energy by replacing hundreds of CPU nodes with a GPU card. This speed-up ratio is also much faster than the speed-up reported by the latest GPU-based methods [10, 29], demonstrating the effectiveness of GWA-X’s regression method and permutation testing strategy.

To evaluate performance scalability on multiple GPUs, GWA-X was run on a system equipped with two NVIDIA A5000 GPU cards. Figure 9 shows GWA-X’s runtimes with 500,000 genetic markers and 430,000 samples, and the number of permutations ranged from 1000 to 5000. For the same number of permutations, the runtime decreases almost linearly with the number of GPUs, and the overhead is only a small fraction (less than 0.82%) of the total runtime. The small overhead demonstrates the effectiveness of overlapping data transfers with computations. These results show that GWA-X’s computational performance is scalable to multiple GPUs.

**Figure 8:**
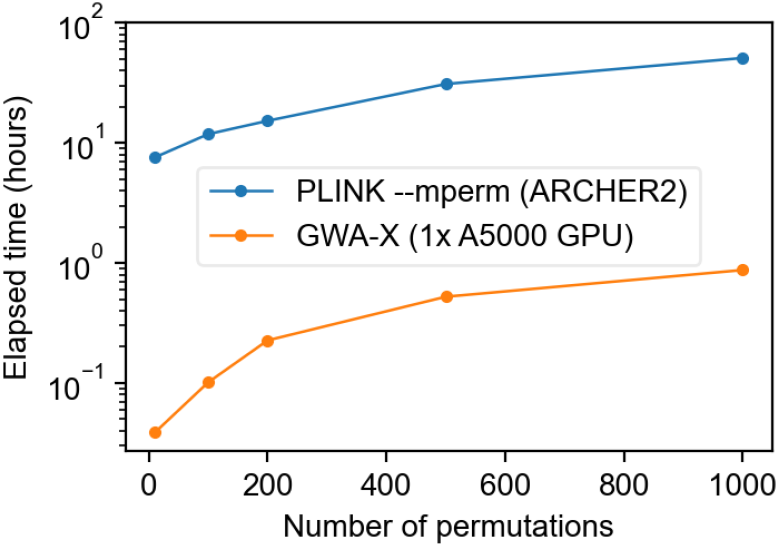
The computational performance of permutation tests with GWA-X and PLINK

**Figure 9:**
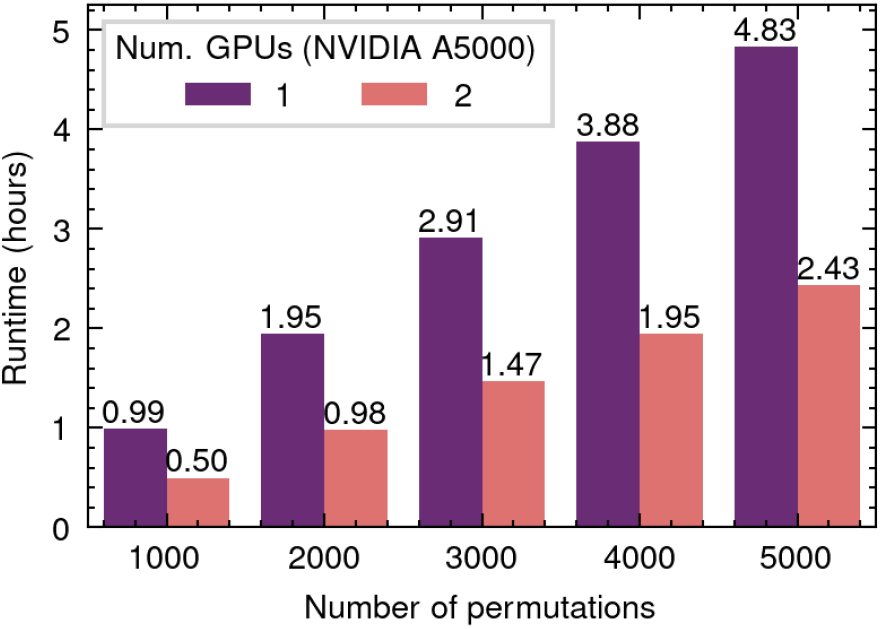
GWA-X’s performance scalability on multiple GPUs

### 6.2 Statistical Performance

GWA-X’s statistical performance is evaluated by comparing the empirical power and false discovery rate (FDR) of the standard genome-wide threshold (5*e−*8) and two permutation-based thresholds at the 5% level obtained from 1,000 and 10,000 permutations (*Q*). For the comparison, 10,000 causal markers (*h*^2^ = 0.2) were simulated on the same dataset used in the computational performance evaluation with the random noise generated from a normal distribution. The significant markers are the markers with *p*-values less than the threshold. The empirical power is the proportion of causal markers detected as significant, and FDR is the proportion of significant markers that are non-causal.

Figure 10 shows the power, FDR, and the number of significant variants detected using the standard and permutation-based significant thresholds. Compared with the standard threshold, the permutation-based thresholds detected more significant variants and achieved higher statistical power while maintaining FDR. The power increased with *Q*, indicating that the accuracy increased with the number of permutations. These results also show the benefit of GWA-X’s acceleration, which allows more permutations to achieve higher accuracy. Previous studies [7, 10] show that GWAS with nonnormally distributed traits are prone to false positive associations. This evaluation compares the statistical performance of permutation-based and Bonferroni-adjusted thresholds on the simulated dataset of ∼1M markers with 5,000 causal markers (*h*^2^ = 0.2) and random noise generated from Student’s t and gamma distributions. The Student’s t-distributed noise (Std-t) was generated with a degree of freedom parameter of 4, and gamma-distributed noise was generated with shape parameters of 1 (gamma-1) and 2 (gamma-2). The permutation-based thresholds were the thresholds at the 5% level from 1,000 permutations.

**Figure 10:**
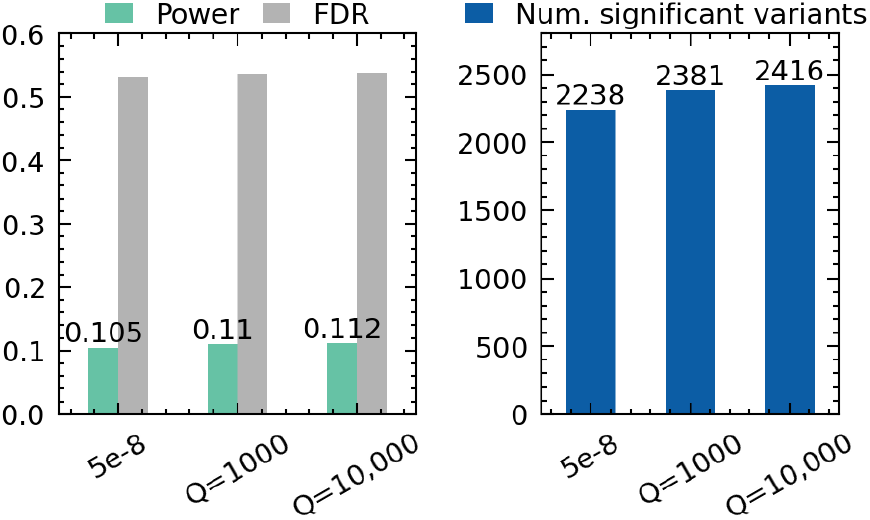
The empirical power, FDR, and number of significant markers of permutation-based thresholds

**Figure 11:**
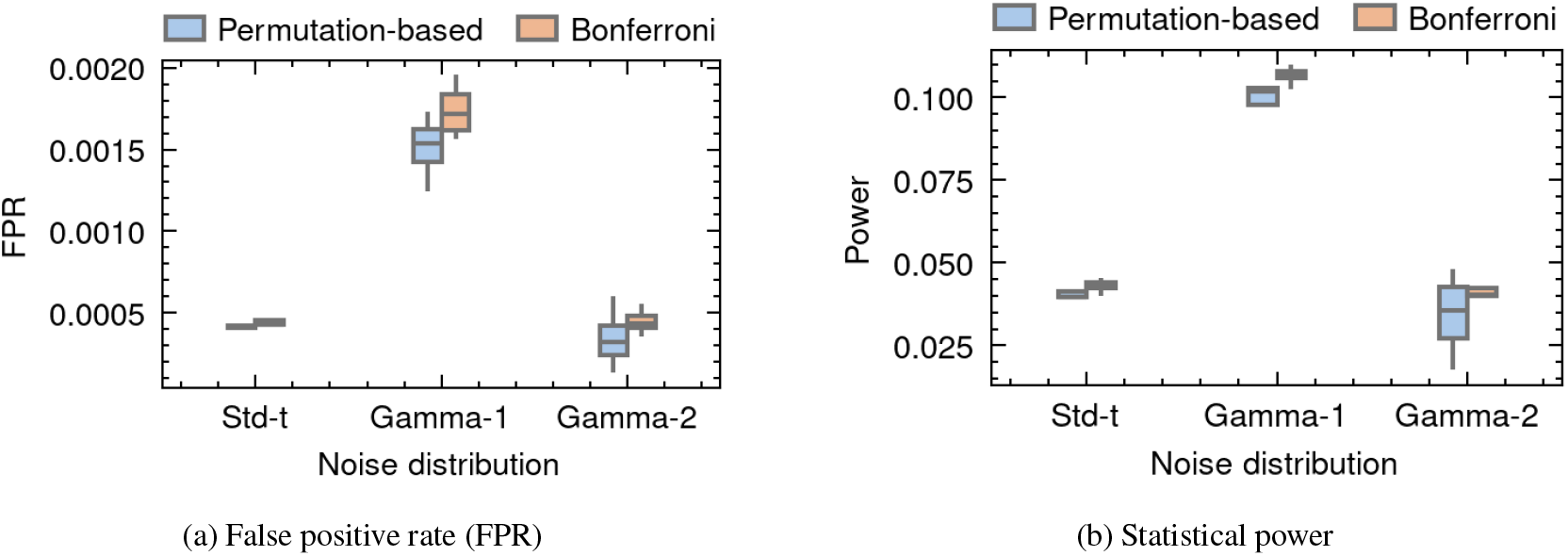
The empirical (a) false positive rate and (b) power results of permutation-based and Bonferroni thresholds on skewed traits. Each box plot represents the distribution of the estimated quantity across 10 simulation replicates.

Figure 10 shows the false positive rate (FPR) and power results of permutation-based and Bonferroni thresholds on the dataset with the non-normal noise distributions. FPR is the proportion of non-causal markers detected as significant. Permutation-based thresholds reduced FPR while maintaining power, with a higher FPR reduction shown in traits with gamma-distributed noise. For the Student’s t-distributed noise, permutation-based thresholds showed marginal FPR reductions over Bonferroni thresholds, indicating that permutation-based thresholds become more stringent when the trait distribution becomes more skewed from normal, as also suggested by other recent work [10]. These results also show the benefit of permutation testing in reducing false positive associations in non-normally distributed traits.

## 7 Discussion and Conclusion

Currently, there is no GPU-accelerated software for GWAS permutation testing that is both efficient and extensible. GWA-X employs a novel whole-genome regression method and a CPU-GPU collaboration strategy to fully utilize GPU computing capabilities. The computational efficiency and extensibility achieved simultaneously by its architecture are essential for emerging data scales and analyses in GWAS. The evaluation demonstrates unprecedented acceleration and energy saving on regression and permutation tests. Thus, GWA-X provides computational efficiency and flexibility for permutation testing in current and future GWAS.

As future work, GWA-X will be extended to support binary trait analysis, adaptive permutation, and ridge regression. In addition, regression models and permutation tests are used in various applications in omics research. Thus, another direction for future work is extending the regression method and the permutation test approach for different analyses, such as rare variant and single-cell RNA sequencing analyses.

## 8 Acknowledgments

This work used the ARCHER2 UK National Supercomputing Service (https://www.archer2.ac.uk). The work is funded by the Biotechnology and Biological Sciences Research Council (BBS/E/D/10002070, BBS/E/D/30002275, BBS/E/RL/230001A), Health Data Research UK (HDR-9003, HDR-9004), the Medical Research Council (MC_UU_00007/10) and the National Institute for Health Research (NIHR202639).

## Footnotes

† https://git.ecdf.ed.ac.uk/magung/gwa-x

